# A new guest to the blood feast: a novel symbiotic lineage associated with a haematophagous leech from the genus *Haementeria*

**DOI:** 10.1101/2023.12.21.572949

**Authors:** Víctor Manuel Sosa-Jiménez, Sebastian Kvist, Alejandro Manzano-Marín, Alejandro Oceguera-Figueroa

## Abstract

Similarly to other strict blood-feeders, leeches from the *Haementeria* genus (Hirudinida: Glossiphoniidae) have established a symbiotic association with bacteria harboured intracellularly in oesophageal bacteriomes. Previous genome sequence analyses of these endosymbionts revealed co-divergence with their hosts, a strong genome reduction, and a simplified metabolism largely dedicated to the production of B vitamins, which are nutrients lacking from a blood diet. *Candidatus* Providencia siddallii has been identified as the obligate nutritional endosymbiont of a monophyletic clade of Mexican and South American *Haementeria* spp. Nonetheless, given a lack of molecular investigations, the identity of the symbiont housed in the bacteriomes of its sister clade of Central and South American congeners remained unknown. In this work, we report on a novel bacterial endosymbiont found in a representative from this *Haementeria* clade. We found that this symbiont lineage has evolved from within the *Pluralibacter* genus, known mainly from clinical, but also environmental strains. Similarly to *Ca*. Providencia siddallii, the *Haementeria*-associated *Pluralibacter* symbiont displays clear signs of genome reduction, accompanied by an A+T-biased sequence composition. Genomic analysis of its metabolic potential revealed a retention of pathways related to B vitamin biosynthesis, supporting its role as a nutritional endosymbiont. Finally, comparative genomics of both *Haementeria* symbiont lineages suggests that the ancient *Providencia* symbiont was likely replaced by the novel *Pluralibacter* one, thus constituting the first reported case of nutritional symbiont replacement in a leech without morphological changes in the bacteriome.

## Introduction

Obligate symbiosis is a widespread phenomenon across a variety of animals, particularly within those with a nutrient-limited diet (e.g. plant sap or blood; Bennett and Moran 2013; Gu *et al*. 2023; Husnik and McCutcheon 2016; Manzano-Marín *et al*. 2023a; Matsuura *et al*. 2018). Once these obligate associates establish a stable vertical transmission, their genomes undergo radical changes, which include loss of dispensable genes/pathways, an invasion by and subsequent loss of mobile elements, and a shift in nucleotide composition (i.e. becoming more commonly A+T biased), among others (Baumann, 2005; Latorre and Manzano-Marín, 2017; McCutcheon and Moran, 2012; McCutcheon *et al*., 2019; Moran *et al*., 2008). This *genome reduction syndrome* can eventually lead to the impairment of essential functions of the symbiotic bacteria due to pseudogenisation or gene loss affecting key pathways to its own maintenance or that of its host. This impairment can lead to one of three outcomes: breakdown of the symbiosis, co-obligate symbiont acquisition (*i*.*e*. a novel symbiont rescuing losses from the older one), or symbiont replacement. The latter can result in the novel symbiotic bacterium establishing a stable association with its host throughout evolutionary time, thus following a similar genome degeneration path as the former (now extinct) symbiont.

One of the best studied groups of animals with a nutrient restricted diet are sap feeders, which face a strong dietary deficiency of essential amino acids and B vitamins. A group that has historically received less attention are haematophagous animals, which are confronted with a strong B-vitamin deficiency. Not surprisingly, similarly to sap feeders, these obligate blood-feeding organisms house obligate symbionts that are able to compensate for their nutrient-deficient diet (Duron *et al*., 2018; Michalkova *et al*., 2014; Nikoh *et al*., 2014). Often, these microorganisms have been found to inhabit intracellularly in specialised host cells referred to as bacteriocytes, which together make up so-called bacteriomes (Boyd *et al*., 2016; Kikuchi and Fukatsu, 2002; Manglicmot *et al*., 2020; Perkins *et al*., 2005), and have evolved small genomes with reduced gene repertoires that convergently retain B vitamin biosynthetic pathways.

Leeches are a monophyletic group of annelids, which are most (in)famous for their blood-feeding habit. Species in the Glossiphoniidae family are proboscis-bearing leeches, whose blood-feeding members have evolved different bacteriome morphologies to host endosymbiotic bacteria, namely belonging to the alpha- and gammaproteobacteria (Kikuchi and Fukatsu, 2002; Manglicmot *et al*., 2020; Perkins *et al*., 2005; Siddall *et al*., 2004). In particular, species of the *Haementeria* genus can be distinguished for having two pairs of globular sacs attached by thin ducts to the esophagus (Manzano-Marín *et al*., 2015; Perkins *et al*., 2005). These bacteriomes harbor the gammaproteobacterial symbiont *Candidatus* Providencia siddallii (hereafter *Pr. siddallii*), which has been shown to hold an small A+T-biased genome preserving complete pathways for B vitamin biosynthesis (Manzano-Marín *et al*., 2023b). These characteristics, along with congruence between the phylogenetic relations among the leech hosts and symbionts, support the obligate and “ancient” nature of *Pr. siddallii* as a nutritional symbiont of *Haementeria* leeches. Nonetheless, *Pr. siddallii* has only ever been explored in a clade of South American and Mexican leech species, leaving the identity of the symbionts of its sister clade of Central and South American *Haementeria* species unknown.

In this work, we describe the serendipitous discovery of a novel nutritional symbiont belonging to the *Pluralibacter* genus in *Haementeria* leeches. Through whole genome sequencing and genome-based metabolic inference, we show this novel symbiont displays clear signs of an obligate nutritional-based association accompanied by continuous vertical transmission. We corroborate that both *Pr. siddallii* and the novel *Pluralibacter* symbiont show convergent retention of biosynthetic pathways for B vitamins, supporting the equivalent roles they play complementing their hosts’ diet.

## Results & Discussion

### Genome and phylogenetic placement of a novel *Hamenteria* symbiont

As a result of whole-genome sequencing of the novel *Haementeria* sp. COZEM-ANN-HIR-001/002 (hereafter *Haementeria* sp.), a full mitochondrial genome was assembled. Using previously published mitochondrial *cox1* gene sequences, we were able to confidently place the yet-undescribed *Haemneteria* sp. within the clade of Central and South American species as sister to the clade made up by *Ha. paraguayensis* and *Ha. lutzi* (supplementary **figure S1**).

The genome assembly of the novel bacterial symbiont sequenced from *Haementeria* sp. resulted in seven partially-overlapping contigs (*i*.*e*. joined by end-sequences but not resolved) adding up 1.18 Mega base-pairs (**Mbp**) with an average coverage of 1030x. A BLASTN web-search using the 16S sequence of the newly identified symbiont *vs*. the NCBI’s non-redundant nucleotide collection suggested a close relationship of it with the *Pluralibacter* genus (Gammaproteobacteria: Enterobacteraceae). The novel symbiont encodes for a total of 846 intact proteins and retains 78 putative pseudogenes, whose products are either interrupted by a premature stop codon, or are eroded to a point where essential domains are missing. When compared to the *Pr. siddallii* symbionts of other *Haementeria* (**table 1**), it becomes evident that both lineages have experienced genome reduction similar to that of other strictly vertically transmitted symbionts of arthropods (Latorre and Manzano-Marín, 2017; Mahmood *et al*., 2023; Manzano-Marín *et al*., 2020; McCutcheon *et al*., 2019; Říhová *et al*., 2023). However, the novel symbiont of *Haementeria* sp. displays a more moderate degree of genome reduction when compared to *Pr. siddallii* symbionts, namely taking into account the G+C content, number of CDSs, RNA genes, and pseudogenes. This observed difference suggests that the symbiotic relation of the novel symbiont and *Haementeria* is younger to that of *Pr. siddallii* and their hosts. This would follow the prediction from comparative genomics that older more established symbionts have a highly degenerated but stable genome (as *Pr. siddallii*), while more recent ones tend to display larger and less degenerated ones (as the novel symbiont does; Latorre and Manzano-Marín 2017; McCutcheon *et al*. 2019).

**Table 1.** General genomic characteristics of *Haementeria* endosymbionts. PSAC, PSOF, and PSLP denote the *Pr. siddallii* strains associated to *Ha. acuecueyetzin, Ha. officinalis*, and *Ha. lopezi*; respectively. **NA**= complete data not available in annotation. **\***= calculated from the coverage of the collapsed assembly of the rRNA operon.

|  | *Pl. lignolyticus* |  | *Pr. rettgeri* | *Pr. siddallii* |  |  |
| --- | --- | --- | --- | --- | --- | --- |
| Bacterial strain | SCF1 | novel symbiont | AR_0082 | PSAC | PSOF | PSLP |
| genome size | 4.81 Mbp | ca. 1.19 Mbp | 844 kbp | 756 kbp | 844 kbp | 741 kbp |
| G+C content | 57.0 % | 43.5 % | 23.9 % | 22.0% | 23.9% | 23.3% |
| CDSs | 4,399 | 846 | 628 | 610 | 628 | 612 |
| rRNAs | 22 | 9* | 3 | 3 | 3 | 3 |
| tRNAs | 82 | 42 | 33 | 33 | 33 | 34 |
| ncRNAs | NA | 10 | 2 | 2 | 2 | 2 |
| pseudogenes | 50 | 71 | 5 | 4 | 5 | 5 |
| Mobile elements | YES | None | YES | None | None | None |

In order to infer the phylogenetic origin of this novel *Haementeria*-associated symbiont, and in light of the possible close relationship to the *Pluralibacter* genus, we performed a phylogenetic reconstruction using the predicted proteomes of representatives from the *Enterobacteraceae* and *Eriwnaceae* (**figure 1** and supplementary **figure S2**). The phylogenetic reconstruction clearly recovered the novel symbiont nested within the *Pluralibacter* genus together with *Pl. lignolyticus* and *Pl. gergoviae*. Similarly, the 16S rRNA gene identity and *Average Nucleotide Identity* values, when compared to the other sequenced representatives of the genus (between 94.03-95.13 and 74.19-79.48, respectively), provide support this symbiont representing a novel species within the *Pluralibacter* genus (Barco *et al*., 2020; Chun *et al*., 2018; Yarza *et al*., 2014). The phylogenetic reconstruction also evidenced a long branch leading to the novel *Pluralibacter* symbiont, a feature of many long-term vertically-transmitted endosymbiont lineages (Bennett and Moran, 2013; Husník *et al*., 2011; Manzano-Marín *et al*., 2020, 2023a; Ř íhová *et al*., 2023).

**Figure 1.**
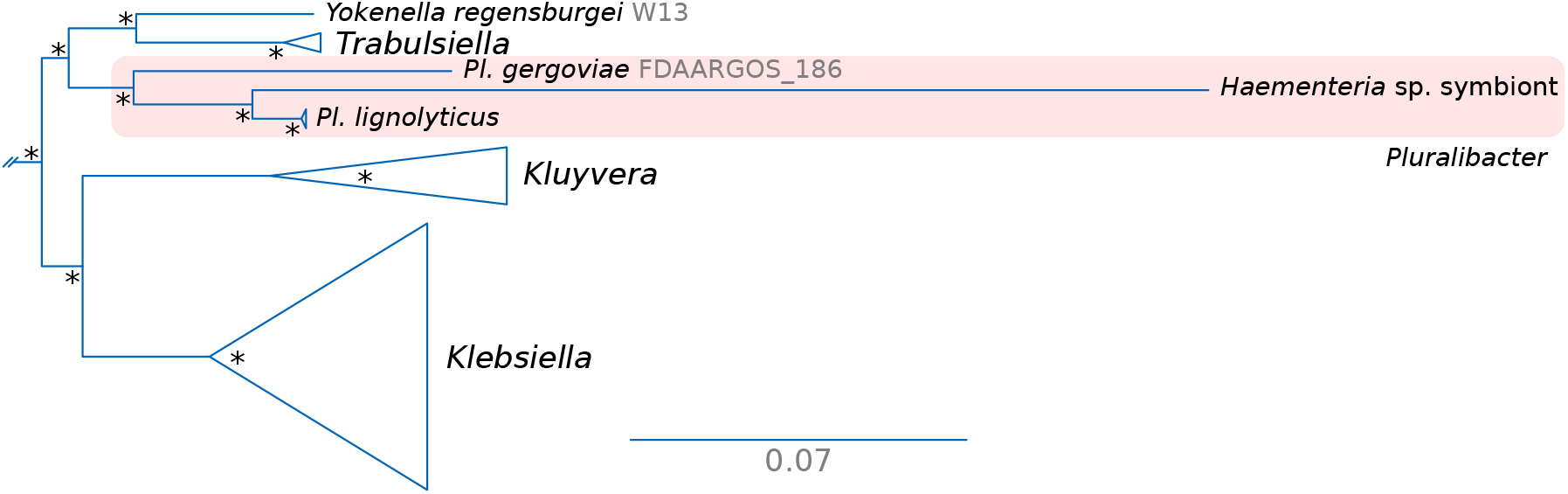
Phylogenetic placement of the novel symbiont of *Haementeria* sp. Excerpt of Maximum-Likelihood phylogenetic reconstruction of *Enterobacteraceae* representatives. The *Pluralibacter* clade is shaded in red. An asterisk at nodes represents an *UltraFast* bootstrap support of 100.

### Metabolic capabilities of the novel *Pluralibacter* symbiont of *Hamenteria* sp

In order to get insight into the genomic reduction and metabolic capabilities of the novel *Pluralibacter* symbiont of *Haementeria* sp, we performed clustering of their protein-coding genes and a COG category analysis to infer the functional profile divergence of the novel symbiont with respect to free-living *Pluralibacter* spp. (supplemetary **figure S3**). Contrasting *Pr. siddallii*, whose genetic repertoire is almost exclusively a subset of that of its free-living relatives (Manzano-Marín *et al*., 2015), the novel *Pluralibacter* symbiont retains a rather large number of strain specific genes (129). This fact suggests that this symbiont likely evolved from within a yet-unknown *Pluralibacter* species, consistent with only two species known within the genus. The two free-living *Pluralibacter* spp. retain a highly similar COG functional profile with some differences in categories *G, T, W*, and *X*; translating to differences in their carbohydrate transport and metabolism, signal transduction mechanisms, extracellular structures, and mobile element content. On the other hand, the novel *Pluralibacter* symbiont shows a distinct profile, with functions such as those involved in cellular maintenance (*J* and *L*), cellular processes and signaling (*D, M*, and *O*), and metabolism (*C, F*, and *H*) overrepresented when compared to the free living *Pluralibacter* spp. This latter category includes genes involved in coenzyme transport and metabolism (*M*), which encompasses all genes involved in B vitamin biosynthesis and which are highly retained in the novel *Pluralibacter* symbiont. This pattern is convergent to that observed in *Pr. siddallii* and other nutritional symbionts from obligate blood feeders (Mahmood *et al*., 2023; Manzano-Marín *et al*., 2015; Říhová *et al*., 2017, 2022, 2023), evidencing a convergent genome reduction among these symbiotic lineages.

The metabolic reconstruction of *Pl. haementericola* (supplementary **figure S4**) indicate that this bacteria can achieve facultative anaerobic respiration, coding for two types of NADH-dehydrogenases (NADH:quinone oxidoreductase I and the Na(+)-translocating NADH-quinone reductase NQR, which could pump H+ and Na+ protons respectively), a fumarate reductase, and a cytochrome bd-I ubiquinol:oxygen oxidoreductase. This is in stark contrast to *Pr. siddallii*, which only preserves genes to perform anaerobic respiration (Manzano-Marín *et al*., 2015). All the main enzymes for the central carbohydrate metabolism are present, thus the endosymbiont would achieve the metabolism of glucose until the production of pyruvate that is then oxidized to obtain acetyl-coA, that then enters to the tricarboxylic acid cycle. Similarly to *Pr. siddallii*, it preserves intact pathways to synthesise all purines and pyrimidines *de novo*.

Regarding the retention of B vitamin biosynthetic genes, which would theoretically compensate for its host’s deficient blood-based diet, the novel *Pluralibacter* symbiont preserves almost-intact pathways for all B vitamin biosynthetic genes (**figure 2**). The one notable exception is the absence of *nudB*. However, the *nudB* gene is not universally conserved across endosymbionts of blood-feeding organisms (Manzano-Marín *et al*., 2015, 2023b; Říhová *et al*., 2023). In fact, it has been shown that many phosphatases display a wide-range substrate specificities in *Escherichia coli* (Haase *et al*., 2013; Kuznetsova *et al*., 2006), suggesting other phosphatase(s) encoded in the symbiont’s genome might be fulfilling a similar role to *nudB*. In contrast to *Pr. siddallii*, the novel *Pluralibacter* symbiont could synthesise NAD from the import of nicotinate, and could still synthesise pantothenate (vitamin B5) from aspartate and valine. In addition, the novel symbiont retains redundant enzymes for some metabolic steps of the B vitamin biosynthetic pathways, unlike *Pr. siddallii* and other well established symbionts (Bennett and Moran, 2015; Koga and Moran, 2014; Manzano-Marín and Latorre, 2016; Manzano-Marín *et al*., 2023b). These metabolic features all support a younger age for the association of *Haementeria* leeches with *Pluralibacter* compared to that with *Provdencia*.

**Figure 2.**
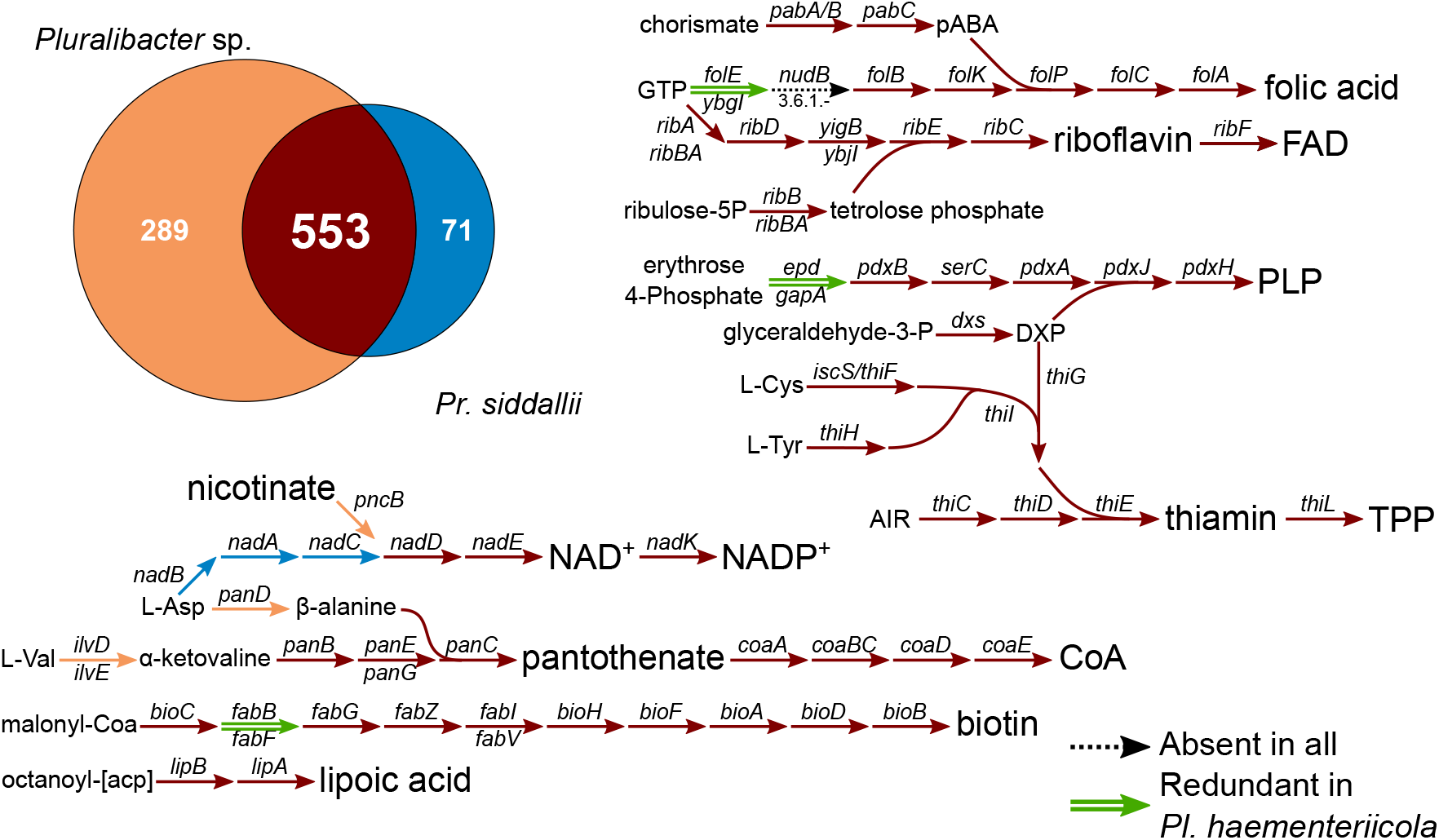
Shared genes and B vitamin pathways of *Haementeria*-associated nutritional symbionts. At the top left, a Venn-like diagram displaying the results of *OrthoMCL* clustering of the predicted proteomes of *Pr. siddallii* from *Ha. officinalis* and the novel *Pluralibacter* symbiont. Diagram of the B-vitamin biosynthetic pathways. Arrows connect metabolites, and names on them indicate the gene coding for the enzyme involved in the enzymatic step.

Regarding amino acids biosynthesis, it has lost the ability to *de novo* synthesise most of the essential amino acids, except for arginine and phenylalanine (from chorismate), as well as non-essential ones (supplementary **figure S4**). However, it retains importers for branched-chain amino acids (isoleucine, leucine, and valine; BrnQ), aspartate (DcuA, DauA), serine and threonine (SstT), and aromatic amino acids (phenylalanine, tryptophan, and tyrosine; AroP). We found evidence that the *Pluralibacter* symbiont was formerly able to code for an methionine-specific ABC transporter complex (MetNIQ), from which a pseudogenised *metN* is retained. Enzymes involved in tryptophan, histidine, and methionine biosynthesis are completely absent. Therefore, the symbiont would largely depend on external sources for the supply of most amino acids. Since after water, proteins are the most abundant organic compound in blood (Lehane, 2010), the host’s diet could represent such an external source to solve both host and endosymbiont essential amino acid requirements.

### ’*Candidatus* Pluralibacter haementeriicola’ *sp. nov*

’*Candidatus* Pluralibacter haementeriicola’ (hae.men.te.ri.i’co.la. N.L. fem. n. *Haementeria*, a leech genus; L. masc./fem. n. suff. -cola, inhabitant, dweller; N.L. masc. n. *haementeriicola*, inhabiting *Haementeria* leeches).

We propose the specific name ‘*Candidatus* Pluralibacter haementeriicola’ for the lineage of enterobacterial endosymbionts from the *Pluralibacter* genus (*Enterobacterales*: *Enterobacteriaceae*) hitherto exclusively found as a bacteriocyte-associated nutritional endosymbiont of *Haementeria* sp. COZEM-ANN-HIR-001/002. Using available genomic data, the closest relative of this symbiont lineage is “*Pluralibacter*(=[*Enterobacter*]) *lignolyticus*”. The only available genome from ‘*Candidatus* Pluralibacter haementeriicola’ displays clear evidence of strong genome reduction and, based on a genome-based metabolic reconstruction, functions as a nutritional endosymbiont providing B vitamins for its leech host. Given the current lack of microscopic imaging of the bacteriomes of its host, its cellular shape and specific localisation within these organs is hitherto unknown.

## Conclusion

Our results provide strong evidence for ’*Ca*. Pluralibacter haementeriicola’ (hereafter *Pl. haementeriicola*) fulfilling the equivalent role to *Pr. siddallii* as a B-vitamin provider for its *Haementeria* leech host. Moreover, the phylogenetic positioning of *Pl. haementeriicola*, its genomic characteristics, and metabolic capabilities suggest an established but younger symbiotic association than that of *Pr. siddallii* with *Haementeria* leeches. While obligate symbiont replacement has been well studied in arthropods (Mao and Bennett, 2020; Sudakaran *et al*., 2017; Šochová *et al*., 2017), other systems have not undergone such a deep study. The aforementioned features from *Pl. haementeriicola*, as well as its host phylogenetic position, suggest that *Pl. haementeriicola* replaced a *Pr. siddallii* symbiont at least in its host species. We expect further information from endosymbionts of leeches from the same Central- and South-american clade of *Haementeria* will clarify whether this novel symbiosis represents a recent symbiont substitution or a more ancient association.

## Materials and Methods

### Leech collection, DNA extraction, and sequencing

*Haementeria* sp. individuals were collected in Ciénega de las Macanas, Provincia de Herrera, Panamá (06.06.2017; 8.10303 N, 80.593974 W) by Sebastian Kvist, Danielle de Carle, and Alejandro Oceguera-Figueroa under permit number SE/A-61-17 (Ministerio de Ambiente de Panamá). Voucher specimens were deposited in the *Colección Zoológica Dr. Eustorgio Méndez* (CoZEM; https://www.gorgas.gob.pa/coleccion-cientifica/) from the *Instituto Conmemorativo Gorgas de Estudios de la Salud* (Panamá) under voucher numbers COZEM-ANN-HIR-001 and COZEM-ANN-HIR-002. Upon expert inspection of these individuals, it was concluded that they putatively represent specimens of a new leech species that remains to be described. The bacteriomes of two individuals were dissected and total DNA was extracted using a commercial extraction kit (*DNeasy Blood & Tissue Kit*, Qiagen, Hilden, Germany), following the manufacturer’s instructions for purification of total DNA from animal tissues. DNA libraries were constructed using the *NGS Nextera FLEX DNA library preparation kit* (*Illumina Inc*., San Diego, CA) according to the manufacturer’s protocol, and total DNA was multiplexed together with 11 other samples and sequenced on a single lane on the *HiSeqX* platform (150 bp paired-end).

### Genome assembly and annotation

Sequencing reads were right-tailed clipped (quality threshold of 20) using FASTX-Toolkit v0.0.14 (http://hannonlab.cshl.edu/fastx_toolkit/, last accessed December 11, 2023). We then removed reads containing undefined nucleotides (”N”) and those shorter than 75 base-pairs (bps) using PRINSEQ v0.20.4 (Schmieder and Edwards, 2011). The resulting filtered reads were then assembled using SPAdes v3.10.1 (Bankevich *et al*., 2012) (-kmer 55,77,99 -only-assembler) and contigs with a coverage lower than 100 were discarded. Surviving contigs were binned using results from a BlastX v2.11.0 (Altschul *et al*., 1997) search (best hit per contig) against a database consisting of the proteome of *Hellobdella robusta* as well as that of a selection of bacterial strains (supplementary **table S1**). This binning resulted in 10 contigs of putative bacterial origin (coverage *≥*1000) and 13 assigned to the mitochondrial bin (coverage *≥*100). The completeness of molecules assigned to both bacteria and mitochondrion was corroborated by both manual inspection of contigs showing a higher coverage than the background (i.e. host contigs) using the BlastX webserver (*vs*. nr) as well as by inspection of assembly graphs. In no case did we find any additional contigs of putative bacterial origin. The aforementioned binned contigs were used for mapping back the reads from *Haementeria* sp. using Bowtie2 v2.5.0 (Langmead and Salzberg, 2012). This was followed by re-assembly of the mapped reads using SPAdes as described above. This resulted in a single mitochondrial contig and 10 contigs (with overlapping ends) belonging to the putative bacterial endosymbiont.

Draft annotation of the genomes was done using MITOS v6b33f95 (Bernt *et al*., 2013) and Prokka v1.14.16 (Seemann, 2014). This draft annotation was followed by manual curation of the protein-coding genes’ coordinates as well as an update to the product naming using UniProtKB (Bat, 2021), Pfam v34 (Mistry *et al*., 2021) and InterProScan (Jones *et al*., 2014). Annotations for non-coding RNAs was refined using Infernal v1.1.4 (Nawrocki and Eddy, 2013), with the Rfam database v14.1 (Kalvari *et al*., 2021), tRNAScan-SE v2.0.9 (Chan *et al*., 2021) and ARAGORN v1.2.41 (Laslett, 2004). Pseudogenes were manually identified through online BlastX searches of the intergenic regions as well as through BlastP, DELTA-BLAST (Boratyn *et al*., 2012), and domain searches of the predicted open reading frames. Proteins were considered to be putatively functional if all essential domains for the function were found, if a literature search supported the truncated version of the protein as functional in a related organism, or if the predicted protein displayed truncations but retained identifiable domains. Details of the results of these searches are captured in the annotation files. All manual curation was done using UGENE v1.34.0 (Okonechnikov *et al*., 2012).

The fully annotated contigs underwent metabolic annotation in Pathway Tools v22.0 (Karp *et al*., 2021) based on BioCyc and MetaCyc databases (Caspi *et al*., 2020). Contigs were also submitted to the KEGG Automatic Annotation Service (Moriya *et al*., 2007). After automatic reconstruction, manual curation of the databases was done by comparing to known reactions and complexes present both in BioCyc and KEGG. Global metabolic reconstruction was manually edited with Procreate v5.3.6.

### Phylogenetic analysis

For phylogenetic placement of the newly sequenced *Haementeria*-associated endosymbiont, and informed by the 16S identity of it *vs*. available sequences in NCBI’s nr, we downloaded 147 representative genomes from *Enterobacteraceae* and *Erwinaceae*. Next, OrthoFinder v2.5.4 (Emms and Kelly, 2019) was used to group predicted protein coding genes across selected genomes in families of orthologous proteins. We then those consisting exclusively of single-copy core genes (i.e. present across all genomes), which were then individually aligned using MAFFT v7.490 (-maxiterate 1000 -localpair; Katoh and Standley 2013) and fed to Gblocks v0.91b (Talavera and Castresana, 2007) for removal of divergent and ambiguously aligned blocks. The resulting alignment were then concatenated and phylogenetic inference was done using IQtree v1.6.12 (LG4X+I; Minh *et al*. 2020) with 1000 UltraFast bootstrap replicates (Hoang *et al*., 2018). Tree plot was visualised and exported for editing using Figtree v1.4.4 (https://github.com/rambaut/figtree, last accessed December 11, 2023).

For phylogenetic placement of *Haementeria* sp., we followed Oceguera-Figueroa (2012) and collected cox1 gene sequences from representatives of *Placobdella, Helobdella*, and *Haementeria* genera. Sequences were aligned using MAFFT as described above. Phylogenetic inference was performed using Mrbayes v3.2.7(GTR+I+G4; Ronquist *et al*. 2012) running two independent analyses with four chains each for up to 1,000,000 generations and checked for convergence (average standard deviation below 0.001) with a burn-in of 25%. Tree visualisation and exporting was done as described above.

### COG assignment and functional profiles

In order to investigate the differences in functional profile between the symbiont of *Haementeria* sp. and its closest free living relatives (”*Pluralibacter*[=*Enterobacter*] *lignolyticus*” strain SCF1 and *Pluralibacter gergoviae* strain FDAARGOS 186), Clusters of Orthologous Groups categories were assigned to all predicted CDSs with the WebMGA web server (Wu *et al*., 9 07). For assessing functional divergence of the novel leech symbiont from their free-living relative strains, a mean of per COG category frequency was calculated for the latter and subtracted from the given category for both the leech symbiont and *Pluralibacter* strains as in (Manzano-Marín *et al*., 2015). The results were visualized by means of a heatmap plotted in R (R Core Team, 2021) using the function *pheatmap*.

All figures were edited and exported using Inkscape v1.1.2 (https://inkscape.org, last accessed December 11, 2023), unless otherwise stated.

## Supplementary Material & Data Availability

Supplementary **figures S1-S4** and **table S1** have been included in this submission. All auxiliary files and data for other analyses as well as the annotated genome of *Pl. haementeriicola* are available online at https://doi.org/10.5281/zenodo.10091738. Newly sequenced and annotated genomes are in the process of being accessioned at the European Nucleotide Archive (ENA).

## Acknowledgements

This paper is part of the requirements for obtaining a Doctoral degree at the *Posgrado en Ciencias Biológicas, UNAM* of VMS-J. This project was funded by grants from *PAPIIT-UNAM-IN213520* to AO-F, *NSERC* to S.K., and a *Peer Review Grant* from the *Royal Ontario Museum* to S.K. We thank the *Posgrado en Ciencias Biologicas, UNAM* and *CONACyT* for providing a scholarship to VMS-J. AMM has received funding from *European Union*’s *Horizon 2020 Research and Innovation Programme* under the *Marie Sklodowska-Curie* Grant Agreement No. [840270] (LEECHSYMBIO). The authors thank Prof. Dr Thomas Rattei and his team for maintaining the *Life Science Compute Cluster* (LiSC; https://cube.univie.ac.at/lisc) that was used for computational analyses. The funders had no role in study design, data collection and analysis, decision to publish, or preparation of the manuscript.

