## Supplementary figures S1-4 for "A new guest to the blood feast: a novel symbiotic lineage associated with a haematophagous leech from the genus *Haementeria*"

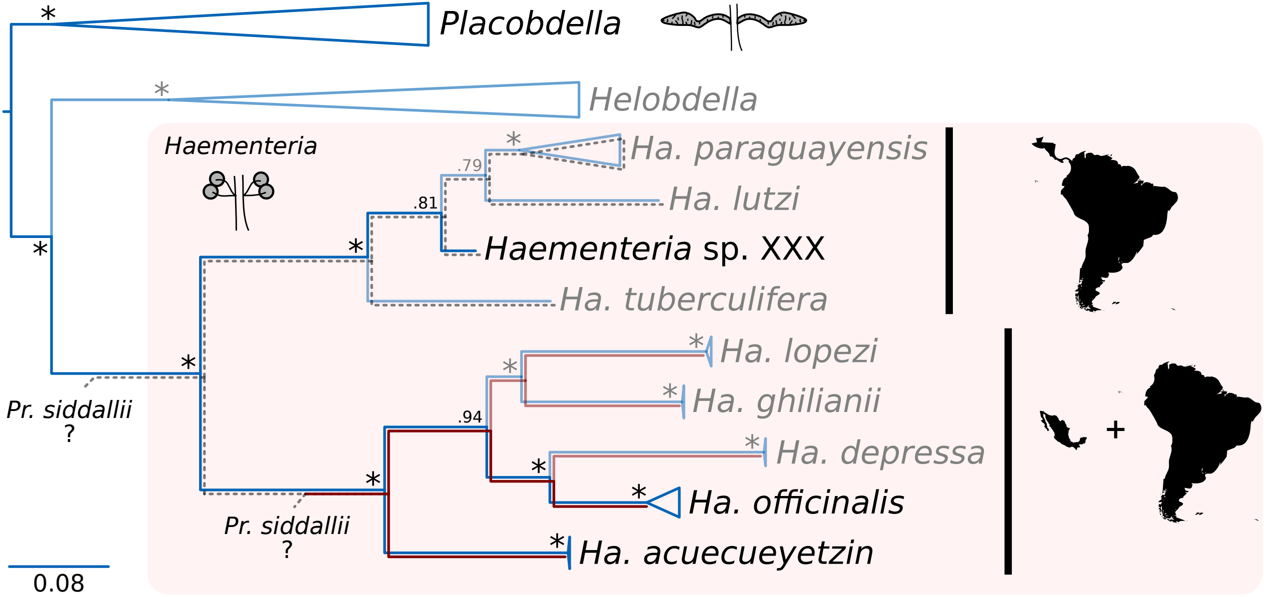


**FIgure S1.** Phylogenetic tree displaying relationships among *Haementeria* species, using *Placobdella* spp. and *Helobedella* spp. as outgrous. Diagrams of the bacteriomes of *Haementeria* and *Placobdella* are shown. Red lines indicate the known presence of the *Pr. siddallii* symbiont and its co-divergence with *Haementeria* hosts. Grey dotted lines indicate the possible (but unkown) presence of a nutritional symbiont, *Pr. siddallii* or otherwise. "?" indicates uncertainty about the identity of the infection and establishment of *Pr. siddallii* as the obligate nutritional symbiont of *Haementeria* spp. Numbers at node represent bootrap support. Asterisks at node represent a bootstrap support of 100%.

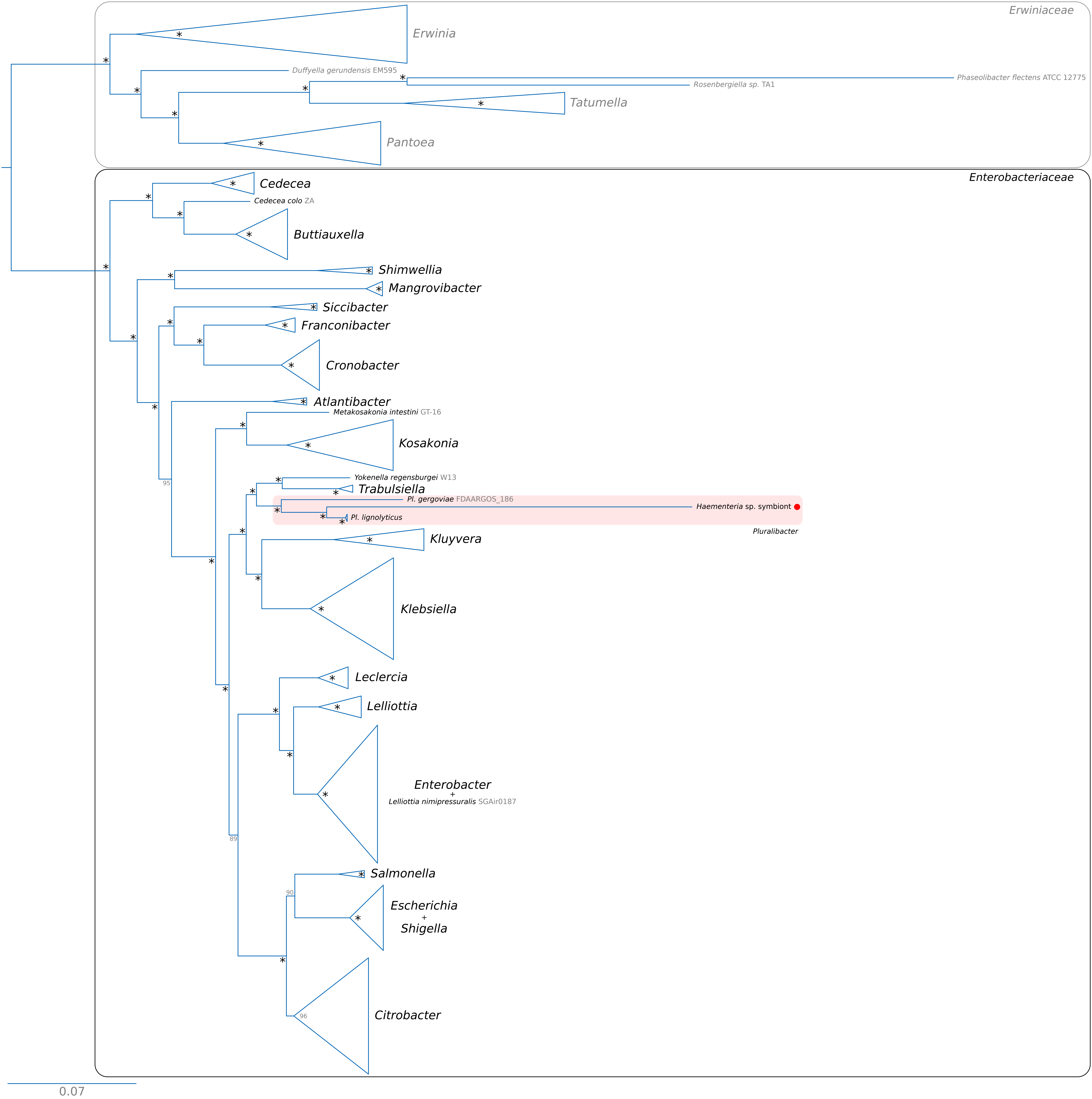


**FIgure S2.** Full collapsed phylogenetic tree displaying relationships among selected *Erwiniaceae* and *Enterobacteriaceae*. The *Pluralibacter* clade is shaded in light red, and the positioning of the novel symbiont is marked with a red dot. Numbers at node represent bootstrap support. An asterisk at nodes translates to a bootstrap support of 100%.

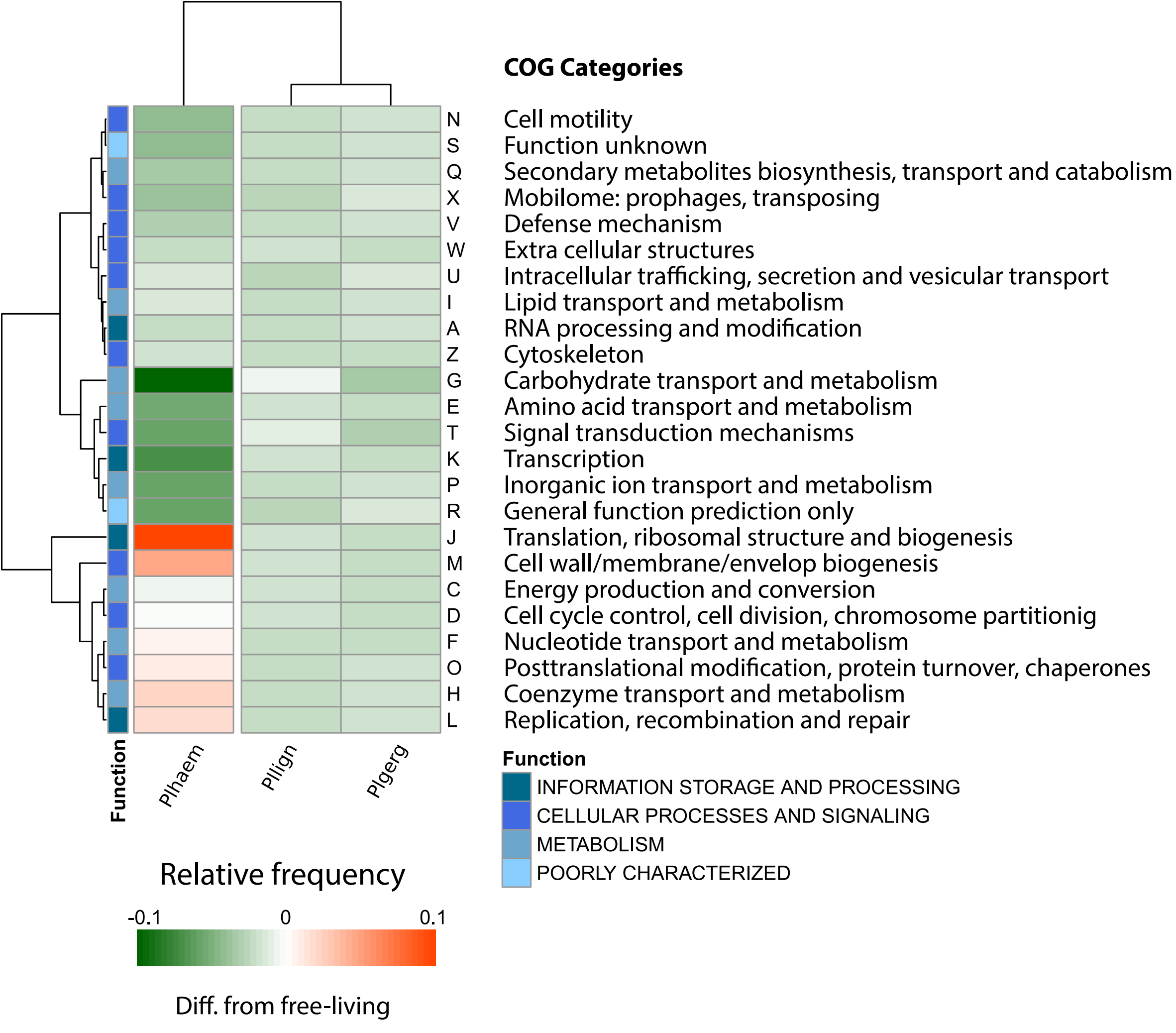


**FIgure S3.** Heatmap displaying a two-way clustering of the functional profile divergence of Pl. haementeriicola compared with free-living Pluralibacter strains. At the bottom, colour key for the difference in relative frequency of each COG category. At the right, corresponding COG categories and colour code for the functional grouping of COG categories."Plhaem"= *Pl. haementeriicola*, "Pllign"= *Pl. lignolyticus*, "Plgerg"= *Pl. gergoviae*.

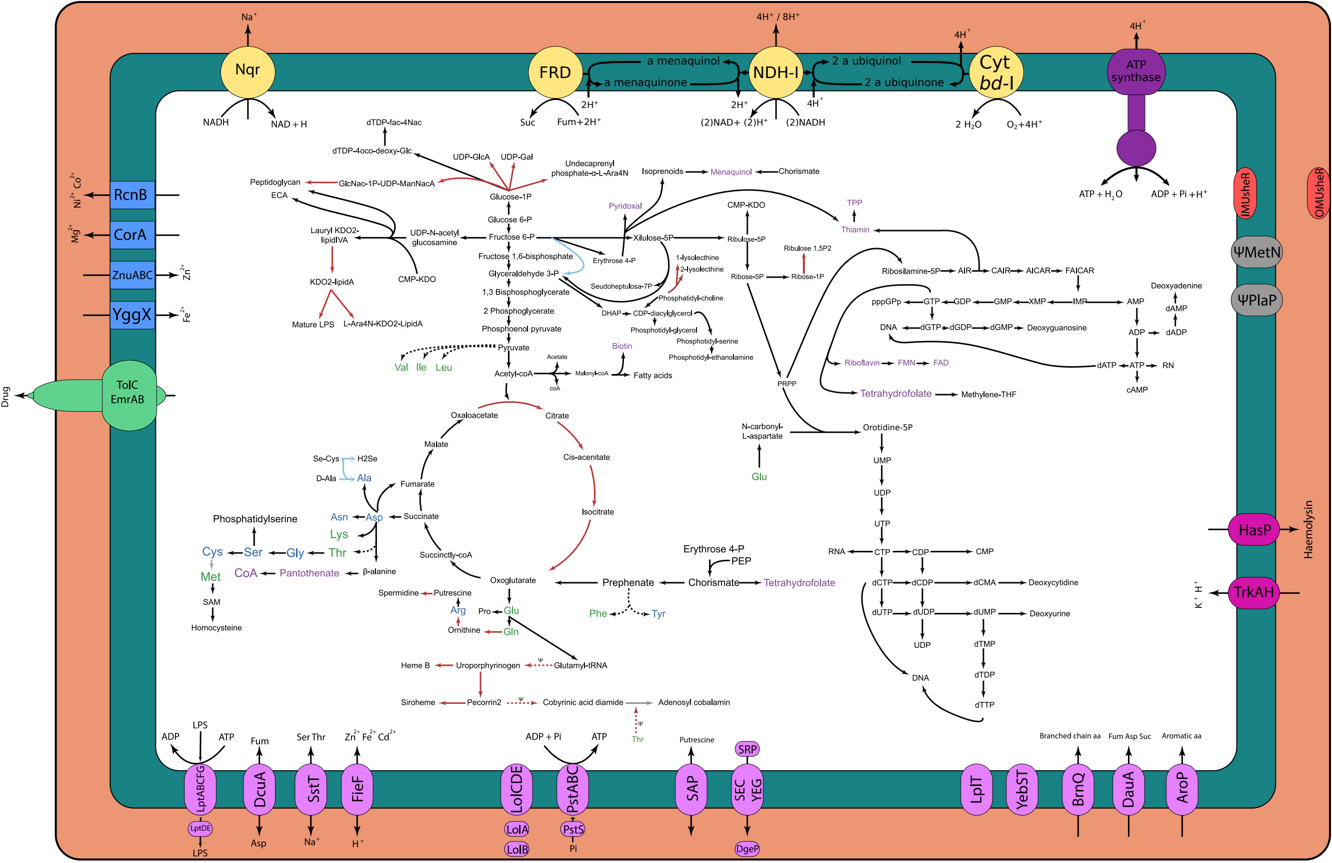


**FIgure S4.** Inferred metabolism of *Pl. haementeriicola* and comparison *vs. Pr. siddallii*. The three coloured compartments represent (from inner to outermost) the cytoplasm, inner membrane, and periplasm. The different membrane transporters are represented by coloured items on their corresponding compartment. Essential and non-essential amino acids are coloured in green and blue, respectively. Coenzymes are coloured in purple. Lines connecting compounds represent enzymatic steps. Red lines represent those steps only found in *Pl. haementeriicola*, while blue lines represent those retained in *Pr. siddallii* but not the novel *Pluralibacter* symbiont. Black, grey, and dotted lines represent those steps retained in both *Haementeria*-associated symbionts, absent in both, and those pathways with some missing steps "ψ"= pseudogenised steps.
